# Kinase-gated coincidence detection controls kinesin-driven lysosome transport

**DOI:** 10.64898/2026.03.27.714776

**Authors:** Laura O’Regan, Sarah A Bristow, Zuriñe Antón, Anna Bodzęta, Jessica A Cross, Derek N Woolfson, Mark. P Dodding

## Abstract

Kinesin motors drive long-range intracellular transport through coordinated cargo recognition and conformational autoregulation, yet the mechanisms that selectively control cargo engagement remain unclear. Here, we identify a phosphorylation-dependent gate on kinesin-1 activity mediated by the carboxy-terminal domain (CTD) of kinesin light chain 2 (KLC2). The KLC2 CTD is constitutively phosphorylated on multiple serine residues, suppressing membrane association via its amphipathic helix. We identify the NIMA-related kinase NEK10 as a KLC2-selective regulator of kinesin-1, restraining motor activation, cargo engagement, and lysosome transport; conversely, loss of NEK10 increases membrane association and, together with low-affinity adaptor interactions, promotes lysosome motility. These findings reveal a phosphorylation-regulated protein-lipid coincidence-detection mechanism - a kinesin-kinase code - that integrates adaptor binding with membrane cues to control kinesin-1-mediated transport and provide a mechanistic basis for understanding paralogue and isoform diversity in the kinesin-1 family.

## Introduction

Kinesin-1 family motors drive long-range intracellular transport along microtubules, moving a wide array of cargoes including membrane-bound organelles, ribonucleoprotein complexes, and invading pathogens (*1–3*). How selective engagement of this wide breadth of cargo is achieved remains poorly understood.

Kinesin-1 complexes are heterotetramers of two kinesin heavy chains (KHCs; KIF5A–C paralogues) and two kinesin light chains (KLCs; KLC1–4 paralogues) (*4*). KLCs integrate cargo adaptor protein binding with direct recognition of organelle membranes (*2, 5, 6*). Adaptor proteins carry conserved sequence and structural features including specific short linear motifs (SLiMs) or coiled-coil domains that bind to the KLC tetratricopeptide repeat (TPR) domains and relieve autoinhibition (*1, 2, 5, 7–11*) through a mechanism that enables cooperativity with microtubule bound MAP7 (*12–15*). Direct membrane association is mediated by the carboxy-terminal domain of KLCs (KLC-CTD)(*6*) **(Fig. 1A).**

**Figure 1.**
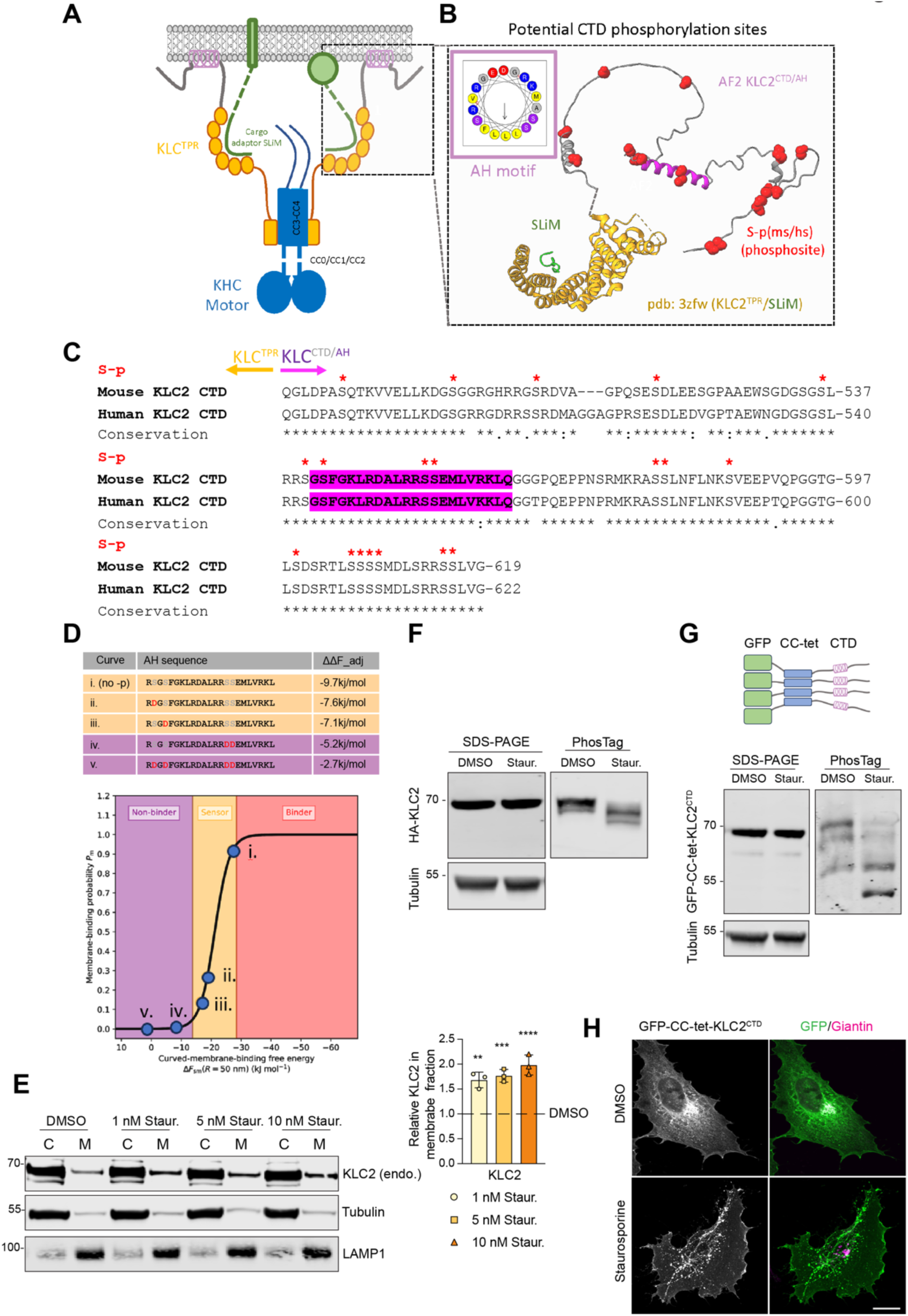
The C-terminal domain of KLC2 is phosphorylated to negatively regulate its association with membranes. (A) Schematic illustrating some general principles of KLC–cargo recognition: adaptor proteins, which may be integral or peripheral membrane proteins, bind to the KLC TPR domains. The mostly unstructured C-terminal domain (CTD) contains an amphipathic helix (AH) that can directly interact with membranes. (B) Integrated structural model showing: (i) recognition of a W-acidic class adaptor SLiM (green) by the KLC TPR domains (orange, pdb:3zfw), alongside an AlphaFold2 model of the KLC2 CTD highlighting the amphipathic helix and annotated with serine phosphorylation sites (red, space fill) curated from the PhosphoSitePlus database. The inset shows a HeliQuest representation of the AH, highlighting its polar and hydrophobic faces. (C) ClustalW sequence alignment of mouse and human KLC2 CTDs highlighting the position of the amphipathic helix alongside curated serine phosphorylation sites from the PhosphoSitePlus database. (D) Results of *in silico* analysis using PMIpred showing predicted membrane-binding properties of the KLC2 AH sequence, compared with those obtained for single and combined phosphomimetic mutations. ΔΔF_adj: the predicted curvature-sensing free energy, corrected for charge effects; ΔF_sm(R=50): the predicted membrane-binding free energy for a vesicle with a 50 nm radius. (E) Western blot and quantification (n = 3) showing results of HeLa cell fractionation and localisation of endogenous KLC2 following treatment with the indicated concentrations of staurosporine (St.). C, cytoplasmic fraction, M, Membrane faction. Statistical significance was determined using one-way ANOVA with Dunnett’s multiple comparisons test. **, p < 0.01, ***, p < 0.001, ****, p < 0.0001. (F) Western blot showing the mobility of HA-KLC2 in HeLa cell extracts treated with vehicle control (DMSO) or 10 nM staurosporine (Staur.) for 1 h. The left panel shows standard SDS–PAGE, and the right panel shows increased mobility on a Phos-tag gel. (G) Schematic showing the design of the GFP-ccTET-KLC2 CTD construct. Western blots show the impact of staurosporine (Staur.) treatment on standard SDS–PAGE and Phos-tag gels. (H) Fluorescence images of Hela cells transfected with GFP-ccTET-KLC2-CTD and stained for the Golgi marker giantin, comparing vehicle control (DMSO) and 10 nM staurosporine (Staur.) treatment for 1 h. Scale bar 10 μm. All data are representative of at least three independent experiments.

The KLC-CTD contains a membrane-induced amphipathic helix (AH) embedded within a predicted intrinsically disordered, basic, serine-rich sequence, which preferentially binds to curved membranes containing negatively charged lipids including phosphatidic acid and phosphoinositides (*6*). This domain is the most prominent site of divergence among the four mammalian KLC paralogues and is further diversified by extensive alternative splicing of KLC1 in vertebrates producing many different long-or short KLCs that, respectively, incorporate or lack the AH and modify its flanking sequences. This suggests that KLC-CTDs function as tunable regulatory modules that shape how kinesin-1 engages distinct cargoes under different cellular contexts (*6, 16, 17*). Consistent with this, altered composition of the KLC pool has been linked to different human diseases, including Alzheimer’s disease associated with a specific KLC1 splice isoform (KLC1E) (*18*), missense mutations in KLC1 associated with schizophrenia (*19, 20*), spastic paraplegia-optic atrophy-neuropathy (SPOAN) syndrome caused by KLC2 upregulation (*21*), and hereditary spastic paraplegia resulting from truncation of KLC4 (*22*). This all underscores the importance of understanding core KLC mechanisms alongside paralogue/isoform specific functions and regulation for both normal cell function and kinesinopathy disease mechanisms (*23*).

Here, we investigate how membrane recognition by kinesin light chains is selectively integrated with adaptor-mediated activation to control kinesin-1 transport. Focusing on KLC2, the ubiquitously expressed light chain and a key determinant of lysosome motility (*6, 24–26*), we identify a phosphorylation-dependent constraint acting through the KLC2 carboxy-terminal domain (CTD). We show that the KLC2-CTD is basally phosphorylated on multiple serine residues, limiting membrane association mediated by its amphipathic helix, and identify the NIMA-related kinase NEK10 as a KLC2- selective regulator of this post-translational modification. Our findings support a model in which phosphorylation restricts coincidence detection between adaptor binding and membrane cues to constrain kinesin-1 activity and organelle transport.

## Results and discussion

### Phosphorylation of the CTD of KLC2 negatively regulates its association with membranes

To begin, we examined KLC2 phosphorylation using the PhosphoSitePlus database (*27*) and mapped curated sites onto an AlphaFold2 (*28, 29*) model and mouse–human sequence alignment of the KLC2 CTD **(Fig. 1B,C & Fig. S1A).** Numerous phosphorylation sites were distributed across the CTD, including several within or adjacent to the amphipathic helix motif. To assess the potential functional impact of these sites, we used PMIpred, a neural network trained on molecular dynamics data (*30*), to model phosphomimetic (serine to aspartate) substitutions that introduce local negative charge, focusing on the helix **(Fig. 1D).** Confirming the utility of this approach, the amphipathic helix was predicted to bind negatively charged but not neutral membranes in a curvature-sensitive manner, consistent with our prior experiments *in vitro* (*6*). Phosphomimetic substitutions at single sites strongly reduced, while combined mutations effectively eliminated the interaction.

To test these predictions experimentally, we examined how kinase inhibition affects endogenous KLC2 localization. Fractionation of HeLa cells treated with low doses of the broad-spectrum kinase inhibitor staurosporine revealed a dose-dependent enrichment of KLC2 in membrane fractions that included the late endosome/lysosomal protein LAMP1 **(Fig. 1E).** PhosTag gel analysis showed a corresponding shift to faster electrophoretic mobility, consistent with reduced phosphorylation, which was not resolved by standard SDS-PAGE **(Fig. 1F).**

To determine whether this effect mapped to the CTD, we analysed a *de novo* designed probe (GFP-CC-Tet-KLC2^CTD^). This construct isolates CTD from other components of the kinesin-1 complex (to uncouple confounding effects of TPR domain adaptor binding) and oligomerises the CTD to enhance membrane avidity (*6*). As for the full-length protein, this construct similarly shifted to faster-migrating, multiple PhosTag-gel specific species upon kinase inhibition, consistent with multisite phosphorylation. Under basal conditions the probe was largely cytosolic, with modest perinuclear enrichment near the Golgi (stained with Giantin) **(Fig. 1G).** Following staurosporine treatment, it redistributed to punctate, tubular and peripheral membranes, indicating that CTD phosphorylation potently suppresses membrane association in cells **(Fig. 1H).**

### NEK10 controls KLC2 phosphorylation

To identify the kinase responsible for this modification, we analysed the sequence of KLC2 CTD using MIT Scansite 4.0 (*31*), alongside the CTDs of KLC1 (Isoform E), KLC3, and KLC4, all of which have an extended CTD with similar amphipathic helices (*6*). Medium-stringency analysis identified consensus phosphorylation motifs for multiple kinases, with additional diversity apparent at low stringency **(Fig. S2).** Notably, members of the NEK family occurred frequently, but differentially between the paralogues, driven by multiple instances of the core [L/M/F/W]-X-X-[S/T] consensus alongside additional NEK family member specific features (*32*).

Based on this enrichment in NEK-family substrate sequences, we performed an siRNA screen targeting all 11 family members using the GFP-cc-Tet-KLC2^CTD^ probe as readout. Cells were scored for staurosporine-like punctate or reticulate membrane association. Depletion of the dual specificity serine/tyrosine kinase NEK10 (*32*) produced a strong and reproducible phenotype, confirmed with two independent siRNAs that achieved >90% NEK10 knockdown **(Fig. 2A–C; Fig. S3A).** NEK10 (and its *C. Elegans* orthologue *nek4l*) has been implicated in many different cellular processes including ciliogenesis, cell-cycle regulation, mitochondrial function, beta-catenin turnover and DNA damage response via p53 (*32–38*). However, to our knowledge, NEK10 has not been implicated in KLC regulation.

**Figure 2.**
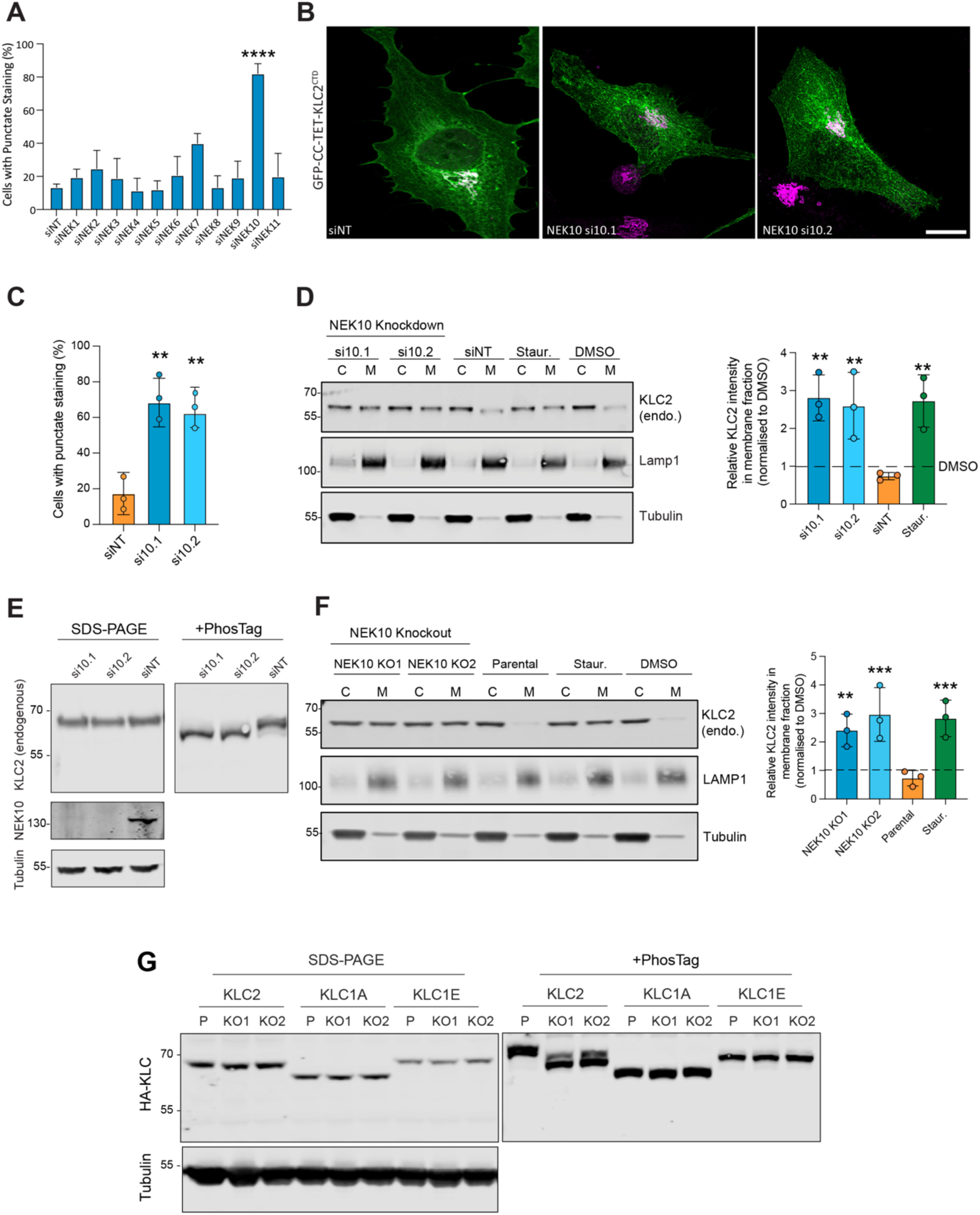
NEK10 controls KLC2 phosphorylation. (A) Results of an NEK-family siRNA screen using pooled oligonucleotides targeting each family member. Cells were transfected with the indicated siRNAs, followed by expression of GFP–CC–tet– KLC2^CTD^, and scored for punctate/reticulate localization (resembling staurosporine-treated cells) or predominantly diffuse localization (resembling control cells). A non-targeting siRNA pool (siNT) was used as a negative control. Cells were fixed and imaged at 72 h post-siRNA transfection. Data represent the mean (± SD) of three independent experiments. Statistical significance was determined using one-way ANOVA with Dunnett’s multiple comparisons test. **, p < 0.01, ***, p < 0.001, ****, p < 0.0001 ****, p < 0.0001. (B, C) Effect of NEK10 knockdown on GFP–CC–tet–KLC2^CTD^ localization using two independent single siRNA oligonucleotides (si10.1 and si10.2). A non-targeting siRNA (siNT) was used as a negative control. **, p < 0.01. Scale Bar is 10 μm (D) Western blot and quantification (n = 3) showing HeLa cell fractionation and localization of endogenous KLC2 following siRNA-mediated depletion of NEK10. C, cytoplasmic fraction, M, Membrane faction. A non-targeting siRNA (siNT) was used as a RNAi negative control. Statistical significance was determined using one-way ANOVA with Dunnett’s multiple comparisons test. **, p < 0.01. (E) Western blot analysis showing the electrophoretic mobility of HA– KLC2 in HeLa cell extracts following NEK10 knockdown, analyzed by standard SDS–PAGE and PhosTag gels. siNT, non-targeting siRNA. (F) Western blot and quantification (n = 3) showing HeLa cell fractionation and localization of endogenous KLC2 in two independent NEK10 CRISPR–Cas9 knockout clones (NEK10 KO1 and NEK10 KO2). C, cytoplasmic fraction, M, Membrane faction. Statistical significance was determined using one-way ANOVA with Dunnett’s multiple comparisons test. **, p < 0.01, ***, p < 0.001. (G) Western blot analysis comparing the electrophoretic mobility of the indicated HA–KLC isoforms in Parental (P) and NEK10 KO HeLa cell extracts analyzed by standard SDS–PAGE and PhosTag gels. The NEK10 knockout–associated PhosTag-dependent mobility shift is specific to KLC2.

We tested whether NEK10 regulates endogenous KLC2. Cell fractionation revealed that NEK10 knockdown caused a marked redistribution of full-length KLC2 from cytosolic to membrane fractions **(Fig. 2D)**, accompanied by increased electrophoretic mobility on PhosTag gels, consistent with reduced phosphorylation **(Fig. 2E).** NEK10 was diffusely distributed throughout the cytosol in immunofluorescence experiments and cell fractionation/western blot analysis showed that it is predominantly cytosolic with a minor membrane-associated pool **(Fig. S3B,C).**

As an orthogonal approach, we generated NEK10 knockout cell lines using CRISPR/Cas9. Two independent clones lacking detectable NEK10 protein phenocopied the siRNA depletion, displaying altered KLC2 phosphorylation and redistribution of KLC2 to membrane fractions **(Fig. 2F,G; Fig. S3D).**

Finally, we assessed paralogue specificity. Only KLC2 exhibited a NEK10-dependent mobility shift on PhosTag gels **(Fig. 2G)**. The long CTD KLC1 isoform (KLC1D) that also contains an amphipathic helix (*6*) **(Fig. S1B)** was unaffected, indicating that NEK10 selectively regulates KLC2 CTD phosphorylation. The short KLC1A that lacks an extended CTD was also unaffected. Together, results from kinase inhibition, siRNA-mediated depletion, and CRISPR/Cas9 knockout converge on NEK10 as a kinesin-1 regulator that selectively controls KLC2 CTD phosphorylation to limit its membrane association.

### NEK10 suppresses kinesin-1 activity by limiting holoenzyme activation and cargo engagement

To determine how NEK10 affects activity of the kinesin-1 holoenzyme, we examined the subcellular distribution of kinesin-1 complexes. Parental HeLa cells were transfected with GFP–KLC2 and HA–KIF5C (KHC). Under control conditions, kinesin-1 complexes were largely diffuse and cytosolic, consistent with their predominantly autoinhibited state (*39*). In contrast, NEK10 KO cells showed an elongated morphology, with GFP–KLC2 accumulating at the distal tips of cellular protrusions, a distribution characteristic of enhanced kinesin-1 activation and plus-end–directed transport (*39, 40*) **(Fig. 3A).**

**Figure 3.**
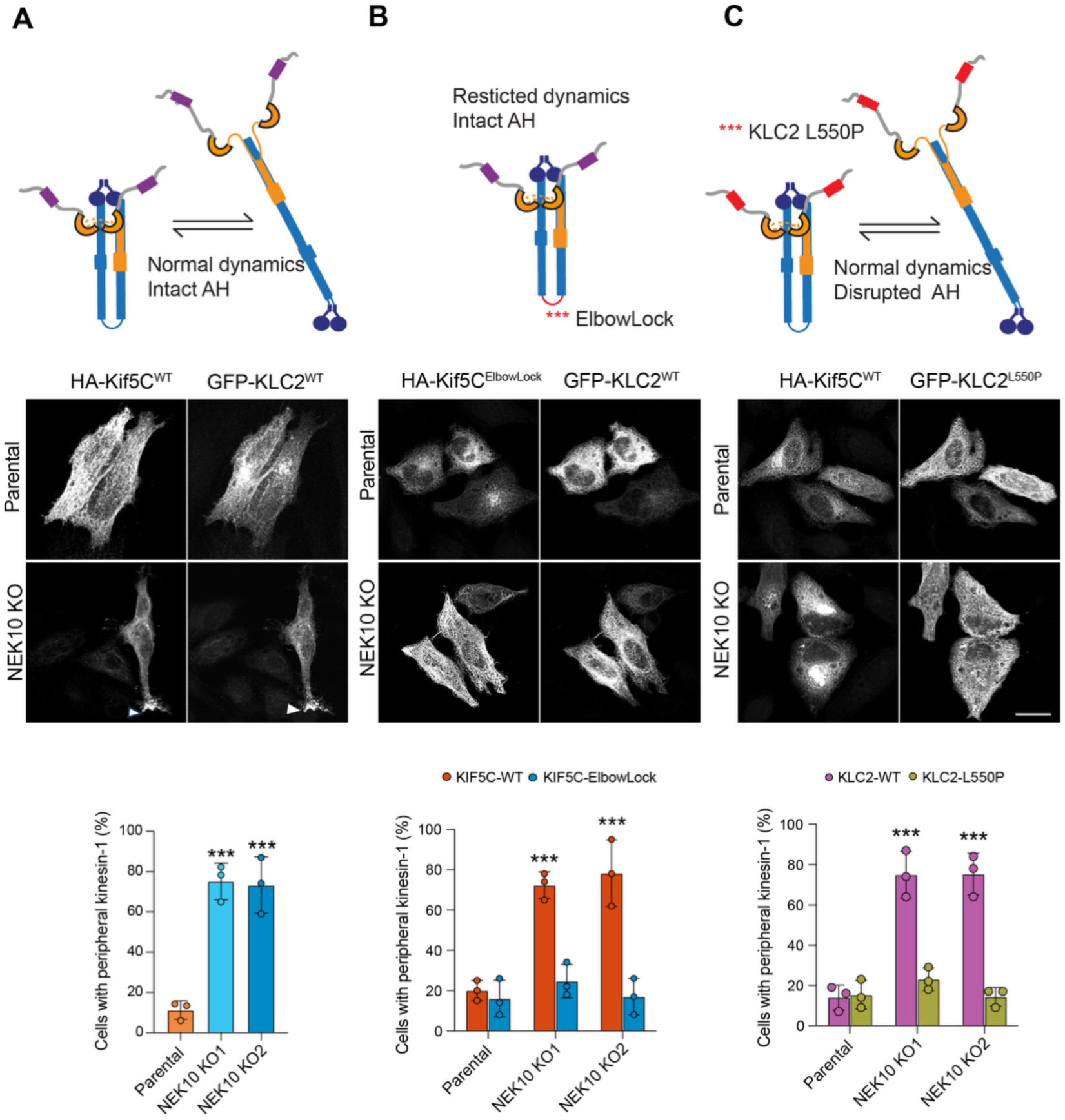
NEK10 suppresses kinesin-1 activity by limiting holoenzyme activation and cargo engagement. Fluorescence microscopy of parental and NEK10 knockout (KO) HeLa cells expressing GFP–KLC2 and HA–KIF5C, used to assess kinesin-1 activity based on accumulation at the cell periphery. (A) Wild-type kinesin-1 complexes are largely diffuse in parental cells but accumulate at distal protrusions in NEK10 KO cells, indicative of enhanced plus-end–directed transport. (B) Expression of the conformationally restricted KIF5C-ElbowLock mutant abolishes peripheral accumulation of GFP–KLC2 in NEK10 KO cells, indicating that NEK10-dependent effects require release from kinesin-1 autoinhibition. (C) Disruption of the KLC2 amphipathic helix (KLC2-L550P) prevents peripheral accumulation in NEK10 KO cells, demonstrating a requirement for AH-mediated membrane engagement. Schematics depict the experimental design and illustrate kinesin-1 conformational activation and KLC2 amphipathic helix function. Scale Bar is 10 μm. Quantification shows the fraction of cells exhibiting peripheral kinesin-1 accumulation. Data represent the mean (± SD) of three independent experiments. Statistical significance was determined using one-way ANOVA with Dunnett’s multiple comparisons test. ***, p < 0.001.

To test whether this redistribution required release from autoinhibition, we co-expressed GFP–KLC2 with a designed KIF5C-ElbowLock mutant, which contains a short deletion in the coiled-coil stalk that prevents the transition from the compact inhibited conformation to the extended active state (*40*). Expression of KIF5C-ElbowLock abolished the peripheral accumulation of kinesin-1 in NEK10 KO cells, indicating that enhanced activity upon loss of NEK10 requires conformational change of the kinesin-1 holoenzyme **(Fig. 3B).**

Next, we asked whether direct membrane association via the KLC2 amphipathic helix was required. In NEK10 KO cells, a KLC2 mutant with a proline mutation that disrupts the hydrophobic face of the helix and blocks membrane binding (L550P) (*6*), remained diffusely cytosolic and failed to accumulate at the cell periphery **(Fig. 3C).** Thus, enhanced kinesin-1 activity upon loss of NEK10 depends on both release from autoinhibition and amphipathic helix-mediated cargo membrane engagement.

Together, these data support a model in which NEK10 suppresses kinesin-1 activity by restricting cargo membrane engagement and subsequent conformational activation. Loss of NEK10 therefore permits kinesin-1 activation through enhanced cargo association and coordinated release from autoinhibition.

### NEK10 constrains W-acidic motif–driven, KLC2-dependent, lysosome transport

Several kinesins, together with cytoplasmic dynein, are recruited to lysosomes through multiple shared, and partially redundant pathways involving several different cargo adaptor proteins and small GTPases (*26, 41*). This redundancy complicates efforts to dissect how kinase-dependent motor regulation is coordinated with adaptor-mediated cargo recruitment. To overcome this complexity and isolate the contribution of NEK10-dependent phosphorylation, we employed a sensitized synthetic lysosome transport model in which kinesin recruitment is controlled by a single, tunable adaptor–KLC interaction (*42*). This system takes advantage of high-resolution SLiM adaptor–KLC TPR structures and defined binding affinities to systematically vary motor engagement independently of endogenous recruitment pathways. Cells were transfected with one of three lysosomal reporters: LAMP1–GFP alone; LAMP1– GFP fused to a single low-affinity (micromolar binder) W-acidic motif from the lysosomal cargo adaptor SKIP (LAMP1–SKIP–GFP); or LAMP1–GFP fused to the engineered high-affinity (low nanomolar binder) KLC ligand KinTag (LAMP1–KinTag–GFP). As reported previously (*42*), under control conditions, LAMP1–GFP–labelled lysosomes were enriched in the perinuclear region, LAMP1–SKIP–GFP produced modest lysosome dispersion, and LAMP1–KinTag–GFP drove robust accumulation of lysosomes at peripheral protrusions and cell vertices **(Fig. 4A,C).**

**Figure 4.**
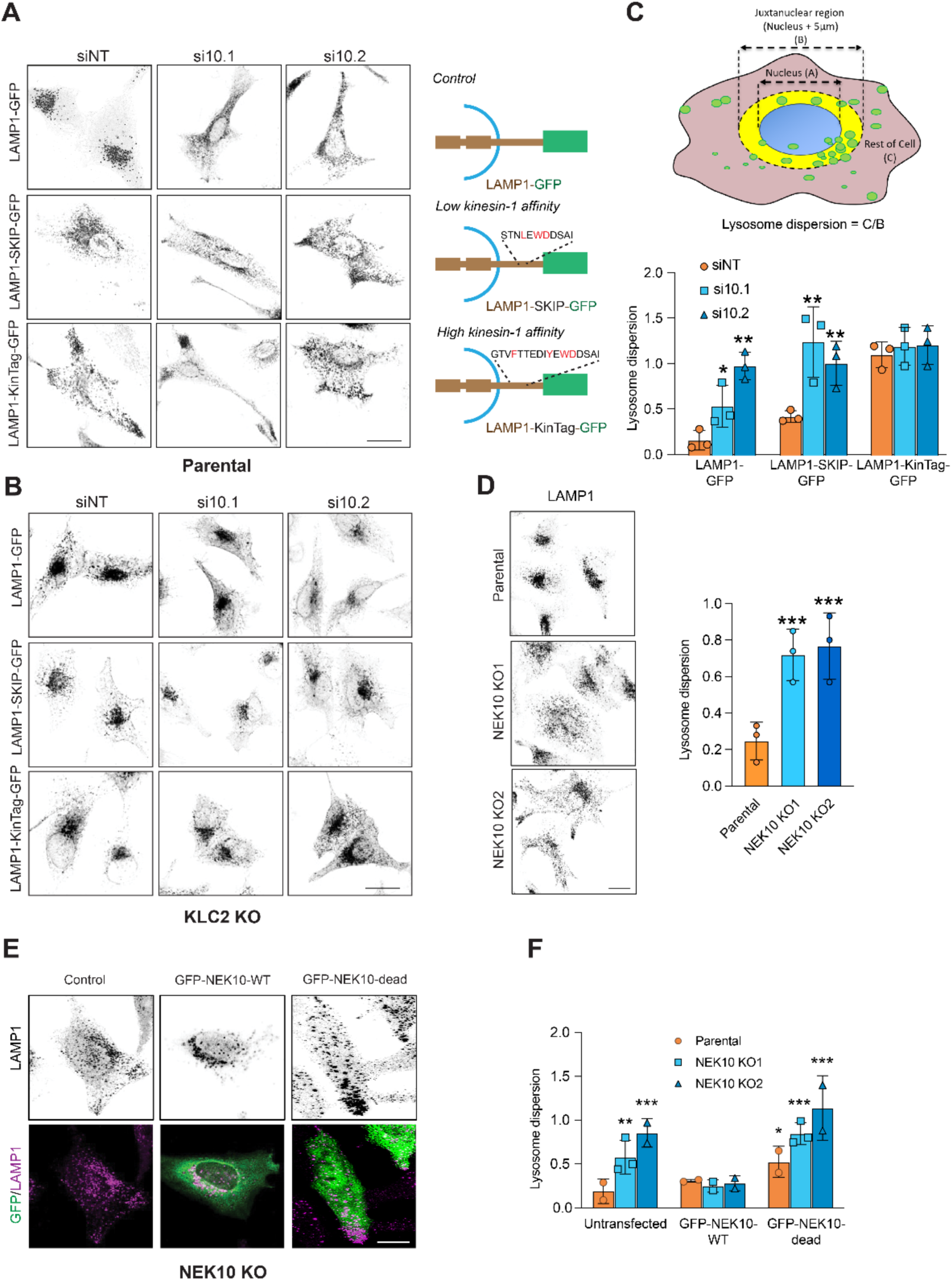
NEK10 constrains W-acidic motif–driven, KLC2-dependent lysosome transport. **(A)** Fluorescence microscopy of (A) parental or HeLa cells or (B) KLC2 KO HeLa cells expressing the indicated LAMP1–GFP reporters: wild-type LAMP1–GFP, LAMP1–SKIP–GFP containing a low-affinity W-acidic motif, or LAMP1–KinTag–GFP containing a high-affinity engineered KLC ligand, with or without siRNA-mediated depletion of NEK10. Schematics illustrate the design of LAMP1 fusion constructs. **(C)** Schematic illustrating quantification approach for lysosome dispersion (*see methods*), and its application for the conditions shown in (A) Data represent the mean (± SD) of three independent experiments. Statistical significance was determined using one-way ANOVA with Dunnett’s multiple comparisons test. *, p < 0.05, **, p, 0.01. siNT, non-targeting siRNA. **(D)** Fluorescence images and quantification (n = 3) showing the distribution of endogenous LAMP1 in parental and NEK10 knockout cells. Statistical significance was determined using one-way ANOVA with Dunnett’s multiple comparisons test. ***, p, 0.001. **(E,F)** Fluorescence images and quantification showing the effects of GFP–NEK10 expression (wild-type or kinase-dead D655N) on lysosome distribution in NEK10 knockout and parental cells. Statistical significance was determined using one-way ANOVA with Dunnett’s multiple comparisons test. *, p < 0.05, **, p < 0.01, ***, p < 0.001. Scale Bar is 10 μm.

These transport phenotypes were strongly dependent on KLC2, as lysosomes remained in the perinuclear region in KLC2 knockout cells (**Fig. 4B)**. In wild-type cells, siRNA-mediated depletion of NEK10 promoted lysosome dispersion in cells expressing LAMP1–GFP, but had no effect in KLC2 knockout cells, consistent with NEK10-dependent regulation of kinesin-1 activity. This effect was further enhanced in cells expressing LAMP1–SKIP–GFP, whereas NEK10 depletion produced no additional dispersion in cells expressing LAMP1–KinTag–GFP **(Fig. 4A–C).**

These results indicate that loss of NEK10 cooperates with natural, low-affinity W-acidic adaptor interactions to promote lysosome transport, but that this regulatory constraint can be bypassed by high-affinity engineered adaptor–motor coupling. Together, the data support a phosphorylation-regulated coincidence-detection mechanism in which NEK10 limits kinesin-1 activation unless adaptor binding is sufficiently strong.

Consistent with the siRNA results, lysosomes were highly dispersed in NEK10 CRISPR-Cas9 knockout cells **(Fig. 4D-E).** This phenotype was rescued by transient expression of wild-type GFP–NEK10, but not by a catalytically inactive mutant (D655N) (*32*) **(Fig. 4E,F).** Moreover, expression of kinase-dead NEK10 in parental cells appeared to act in a dominant-negative manner, inducing lysosome dispersion despite the presence of endogenous NEK10 **(Fig. 4E,F).** Finally, live-cell imaging of NEK10 knockout cells expressing LAMP1-GFP revealed that lysosome dispersion was accompanied by increased lysosome motility **(Fig. S4, Supplementary Movies 1 and 2).**

### NEK10-dependent phosphorylation of the amphipathic helix restricts kinesin-1 activation and membrane association

To identify CTD phosphorylation sites regulated by NEK10, parental and NEK10 knockout (KO) HeLa cells were transfected with constructs expressing full-length N-terminally Citrine-tagged kinesin heavy chain (KIF5C) and N-terminally HA-tagged KLC2. Kinesin-1 complexes were enriched by immunoprecipitation using GFP–TRAP beads, and bound proteins were analysed by mass spectrometry following digestion with either trypsin or endoproteinase AspN to enhance resolution **(Fig. 5A,B).**

**Figure 5.**
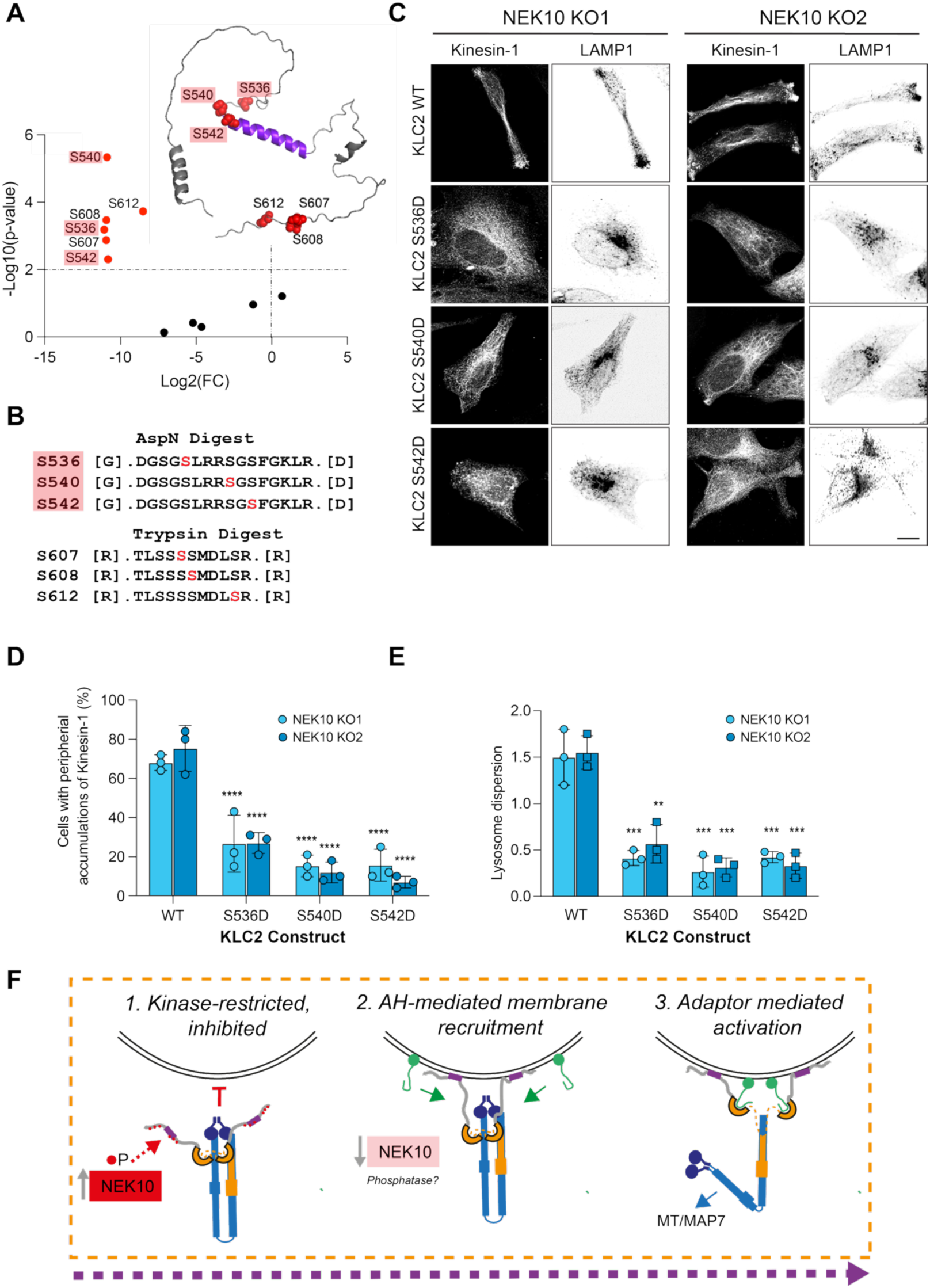
NEK10-dependent phosphorylation of the amphipathic helix restricts kinesin-1 activation and membrane association. **(A,B)** Differential phosphopeptide analysis of KLC2 comparing proteins immuno-isolated from parental HeLa cells and NEK10 knockout (KO) cells. Peptides significantly depleted in NEK10 KO cells (FDR < 0.05; log₂FC < −1) are shown as red dots and mapped onto an AlphaFold2 structural model of the KLC2 CTD, highlighting the amphipathic helix (AH; purple). **(C)** Representative fluorescence images and **(D,E)** quantification of HeLa cells showing the distribution of exogenous kinesin-1 (HA-KIF5C/HA-KLC2) and lysosomes (LAMP1). Wild-type KLC2 complexes accumulate with lysosomes at the cell periphery, whereas this accumulation is suppressed by the indicated phosphomimetic mutations. Data represent the mean (± SD) of three independent experiments. Statistical significance was determined using one-way ANOVA with Dunnett’s multiple comparisons test **, p < 0.01, ***, p < 0.001, ****, p < 0.0001. Scale Bar is 10 μm (F) Integrated model of the kinesin-1 cargo recruitment and activation pathway. Kinesin-1/KLC2 complexes are basally phosphorylated downstream of NEK10, restricting CTD/AH-mediated membrane recruitment. Loss of NEK10 enhances membrane recruitment and promotes subsequent steps in the kinesin-1 activation and transport pathway.

Differential phosphorylation analysis revealed that loss of NEK10 resulted in significantly reduced phosphorylation at six serine residues, all located within the KLC2 CTD. These sites are a subset of phosphorylation events previously reported in phosphoproteomic datasets **(Fig. 1A,B),** with the exception of S612, which, to our knowledge, has not been detected previously. Notably, three residues - S536, S540, and S542 in mouse KLC2 (corresponding to S539, S543, and S545 in the highly conserved human orthologue) **(Fig. 1C),** map within, or immediately adjacent to, the amphipathic helix **(Fig. 5A,B).**

To test whether phosphorylation at these sites is sufficient to regulate kinesin-1 membrane association and activation, parental and NEK10 KO HeLa cells were co-transfected with HA–KIF5C and either wild-type GFP–KLC2 or phosphomimetic GFP–KLC2 variants (serine-to-aspartate substitutions at the identified sites). In NEK10 KO cells, expression of wild-type KLC2 induced the elongated-cell morphology and robust accumulation of kinesin-1 and lysosomes at the cell periphery. In contrast, expression of any phosphomimetic KLC2 variant suppressed these phenotypes, maintaining a diffuse cytosolic kinesin-1 distribution and juxtanuclear lysosome positioning **(Fig. 5C,D,E)**, indicating that acquisition of negative charge at these positions inhibits kinesin-1 membrane association and activation, consistent with the known preference of the CTD for acidic phospholipids (*6*).

### Conclusion

The data presented here demonstrate that phosphorylation of specific serine residues, downstream of NEK10, within the KLC2 carboxy-terminal domain (CTD), including sites proximal to and within the amphipathic helix, is sufficient to impose a constraint on membrane association, kinesin-1 activation, and lysosome transport. This extends understanding of kinesin-1 regulation and lysosome transport beyond adaptor- and MAP7–centric mechanisms (*2, 12, 13, 26*), and provides a direct mechanistic basis for how kinase signalling can gate motor activity and transport **(Fig. 5F).** Viewed more broadly, the kinesin-1 cargo-recognition module functions as a kinase-gated protein–lipid coincidence detector in which phosphorylation modulates how adaptor binding and membrane cues are integrated.

The selectivity of NEK10 for KLC2 over KLC1 suggests that paralogue and isoform diversity within the KLC family enables differential kinase regulation of compositionally distinct kinesin-1 complexes, potentially explaining the expansion and diversification of the KLC family in vertebrates (*6*). Consistent with the roles of NEK kinases in hierarchical signalling networks and substrate priming (40, 41), NEK10 may act within a broader regulatory cascade, perhaps alongside kinases such as AMPK, CKII, and GSK3β previously implicated in KLC regulation (*43, 44*), with opposing phosphatase activities providing further control. Taken alongside established roles for other NEK family kinases in regulation of mitotic kinesins and cilia kinesins (*45–48*) and microtubule dynamics (*38, 49–51*), these findings point to a broader role for NEK signalling in coordinating cytoskeletal organisation and transport. A report of 14-3-3 protein binding to the KLC2-CTD suggests the possibility of an additional regulatory layer (*52*).

Coordinated posttranslational inputs that converge on the CTDs are well suited to tune KLC-cargo recognition in a context-, paralogue-, and isoform-specific manner to specify transport outcomes. In this sense, kinesin-1 regulation may operate through a code-like logic, analogous to the tubulin code, in which combinations of molecular features on tubulin C-terminal tails (CTTs) are interpreted to generate distinct functional outputs (*53*). This framework may provide new opportunities for highly selective intervention in intracellular transport processes that are dysregulated in disease.

## Acknowledgements

This work was funded by BBSRC grants BB/W005581/1 and BB/Z517276/1 to MPD and DNW. JAC was supported by an EPSRC-funded Doctoral Prize Fellowship. MPD acknowledges support from a Lister Institute of Preventative Medicine Fellowship. We are grateful to Dr Kate Heeson and Dr Phil Lewis at the University of Bristol Proteomics Facility for support with phosphopeptide mapping.

## Author contributions

Conceptualization: LO, MPD, DNW; Methodology: LO, SB, AB, JAC, ZA; Investigation: LO, SB, AB, ZA; Formal analysis: LO, SB, AB, MPD; Software / Bioinformatics analysis: MPD; Resources (construct design and generation): JAC, ZA; Funding acquisition: MPD, DNW; Supervision: MPD, DNW; Writing – original draft: LO, MPD; Writing – review & editing: All authors

## Materials and Methods

### Plasmid construction and mutagenesis

HA–KLC2 wild type and the helix-disrupting L550P KLC2 mutant were as previously described (*6*). GFP– CC-tet–KLC2 CTD was generated from de novo–designed tetrameric coiled-coil cDNAs and KLC2 CTD (amino acids 481-619 from NP_001356289.1) synthesized by Eurofins Genomics (pEX-A128 vector) and subcloned into CB6-GFP following a previously described strategy used for variants of KLC1 (*6*). HA-KIF5C-WT and -ElbowLock were as previously described (*39*). LAMP1-GFP, LAMP1-SKIP-GFP, and LAMP1-KinTag-GFP were as previously described (*42*). NEK10 was obtained as a codon-optimized plasmid from Twist Bioscience and subcloned into CB6-GFP using EcoRI/BamHI restriction sites. Point mutations were introduced into GFP-NEK10 and HA-KLC2 using the QuickChange II site directed mutagenesis kit according to the manufacturer’s instructions (Agilent). All constructs were verified by DNA sequencing.

### Bioinformatic analysis of KLC2-CTD

Models of the KLC2 was generated using the AlphaFold2 ColabFold server (*28, 54*) using amino acids 481-619 from NP_001356289.1 as input and prepared for presentation using UCSF Chimera (*55*). Further sequence-structure analysis and ranking was performed using FoldScript (*29*). For phosphorylation site prediction, CTD sequences (following the end of the structurally defined KLC TPR domain (*5*)) to the carboxy terminus of the protein were analysed using MIT Scansite 4.0 for motifs likely to be phosphorylated by specific protein kinases. Sequences used were: KLC2 (NP_001356289.1), KLC1E (NP_001020532.2), KLC3 (NP_803136.2) and KLC4 (NP_001344059.1). The PhosphoSitePlus database (*27*) was queried for KLC2 and outputs show combined data from the human and mouse orthologues. PMIpred (*30*) was run using the peptide settings with negatively charged membrane as target and selected outputs are as presented by the software.

### Antibodies and Reagents

Staurosporine was obtained from Stratech Scientific Ltd. A cell fractionation kit was obtained from Cell Signaling Technology. All siRNAs were obtained from Horizon Discovery. PhosTag SDS-PAGE gels were obtained from Alpha laboratories Ltd. Primary antibodies used were anti-GFP (3E1, Roche), anti-actin (AC-74, Sigma-Aldrich), anti–β-tubulin (AA2, Sigma-Aldrich), anti–histone H3 (Abcam), anti-giantin (PRB-114C, Covance), anti-HA (HA-7, Sigma-Aldrich), anti-KLC2 (ab254848 Abcam), anti-LAMP1 (lysosome-associated membrane protein 1) (D2D11, Cell Signalling Technology), anti-NEK10 (HPA038941, Atlas antibodies). Secondary antibodies were Alexa 568– and Alexa 633–conjugated anti-mouse or anti-rabbit IgGs from Thermo Fisher Scientific.

### Cell culture

HeLa and derived stable cell lines were maintained in Dulbecco’s modified Eagles medium (DMEM) with GlutaMAX-I (Invitrogen) supplemented with 10% heat-inactivated fetal bovine serum (HI-FBS), 100 U/ml penicillin and 100 µg/ml streptomycin, at 37°C in a 5% CO_2_ atmosphere. HeLa cells were obtained from the ATTC, stored in liquid nitrogen and maintained in culture for a maximum of 2 months.

NEK10 KO HeLa cells were generated using CRISPR/Cas9-mediated genome editing. Parental HeLa cells were transfected with the pX459 vector encoding gRNAs targeting NEK10 exon 6 (NEK10 KO1) OR exon 3 (NEK10 KO2) as well as the puromycin resistance cassette. After 48 h, cells were incubated in selective medium supplemented with puromycin for 3-4 d after which cells were diluted for clonal selection into 96-well plates. Following clonal expansion NEK10 KO was verified by Western blot.

Transient transfections were performed using Effectene transfection reagent (QIAGEN) according to the manufacturers’ instructions.

For the NEK siRNA screen, cells were transfected with ON-TARGETplus SmartPool siRNAs (Horizon Discovery) using HiPerFect reagent (Qiagen) according to the manufacturer’s instructions. Final siRNA concentrations were 10 nM and cells were incubated for 72 h before downstream analysis. After identifying NEK10 as a hit, cells were transfected with individual ON-TARGETplus siRNA oligonucleotides targeting NEK10 (J-004052-28, siNEK10.1 and J-004052-29, siNEK10.2, Dharmacon) using the same transfection conditions. Non-targeting siRNA was used as a control in all experiments (D-001810-10, Dharmacon). NEK10 knockdown efficiency was confirmed by immunoblot.

### Protein extraction, immunoblotting and immunoprecipitation

For NEK10 immunoblots cells were lysed into CHAPS lysis buffer (40 mM Hepes (pH7.5), 0.5% CHAPS, 120 mM NaCl, 1 mM EDTA), in all other cases cells were lysed into radioimmunoprecipitation assay (RIPA) buffer (50 mM Hepes (pH 7.5), 150 mM NaCl, 0.1% NP40, 0.5% sodium deoxycholate, 0.1% SDS). Cells were washed in ice-cold 1x PBS prior to lysis in the appropriate ice-cold lysis buffer supplemented with protease and phosphatase inhibitors. Homogenates were incubated on ice for 15 minutes prior to clearing by centrifugation at 12 000xg for 15 minutes at 4°C. Supernatants were collected as soluble fractions, denatured, resolved by SDS-PAGE on NuPAGE 4-12% or Phos-tag (50 µmol/L) SuperSep 7.5% acrylamide precast gels, as indicated, transferred to polyvinylidene fluoride (PVDF) membrane and analyzed by immunoblotting with the antibodies indicated. Western blots were visualized using the Odyssey Infra-red scanning system (LI-COR Biosciences).

Subcellular fractionation was performed using a detergent-based extraction method according to the manufacturer’s instructions, with minor modifications. Briefly, 5 × 10⁶ cells were trypsinized and resuspended in 0.5 ml of 1× PBS. For whole-cell lysates, 100 µl of the cell suspension was mixed with 3× SDS loading buffer containing dithiothreitol (DTT), sonicated (3 × 15 s pulses at 20% power), heated at 95°C for 5 min, and clarified by centrifugation at 15,000 × g for 3 min. The remaining cells were pelleted and resuspended in 500 µl of Cytoplasm Isolation Buffer (CIB) supplemented with protease inhibitors (1×) and PMSF (1 mM), vortexed briefly, incubated on ice for 5 min, and centrifuged at 500 × g for 5 min to obtain the cytoplasmic fraction. The pellet was subsequently extracted with Membrane Isolation Buffer (MIB) to generate the membrane/organelle fraction. All fractions were supplemented with SDS loading buffer containing DTT, heated, clarified by centrifugation, and equal volumes were analyzed by SDS–PAGE. Fraction purity was confirmed by immunoblotting for β-tubulin and LAMP1 which are selectively detected in the cytoplasmic and membrane fractions respectively, with minimal cross-contamination between compartments.

For GFP immunoprecipitations cells were lysed in IP buffer (10 mM Tris/CL (pH 7.5), 150 mM NaCl, 0.5 mM EDTA, 0.5% NP40) supplemented with protease and phosphatase inhibitors, and GFP-tagged proteins immunoprecipitated using GFP-Trap agarose (chromotek) according to the manufacturers’ instructions.

### Immunofluorescence and cell imaging

For fixed cell imaging cells grown on fibronectin-coated coverslips were fixed with 4% paraformaldehyde of 15 min at room temperature or ice-cold methanol for 10 min at −20°C. Cells were blocked with PBS supplemented with 3% BSA and incubated for 30 minutes with primary and secondary antibodies diluted in blocking buffer. DNA was stained with DAPI. Imaging was performed using a Leica SP8 confocal laser scanning microscope (Leica Microsystems, Gernamy) equipped with a 63x/1.4 NA oil immersion objective. Images were acquired using the Leica lightning deconvolution mode with default adaptive settings.

### Quantification of lysosome dispersion and motility

Lysosome dispersion was quantified in Fiji/ImageJ (NIH) using LAMP1 fluorescence. For each cell, a juxtanuclear region was defined as the cytoplasmic area within 5 µm of the nuclear boundary, generated by expanding the nuclear ROI and excluding the nucleus. The remaining cytoplasm was defined as the rest-of-cell ROI. Integrated LAMP1 intensity was measured in both regions, and lysosome dispersion was calculated as the ratio ‘rest-of-cell’ integrated intensity to that in the juxtanuclear region. Identical analysis parameters were applied across conditions.

Live-cell imaging of lysosomes was performed using GFP-tagged LAMP1. Cells were cultured on 35 mm glass-bottomed dishes and imaged using an Olympus iXplore spinning-disk confocal microscope equipped with a SoRa disk using a 60×/1.5 NA oil immersion objective. Single focal plane images were acquired every 240 ms for 2 min. Cells were maintained at 37 °C and 5% CO₂ during imaging, and identical acquisition settings were applied across conditions. Lysosome trajectories were analyzed in Fiji/ImageJ using the TrackMate plugin to quantify lysosome speed and distance.

### Western blot quantification

Western blot quantification was performed on lysates prepared as described above. Equal amounts of protein were resolved by SDS–PAGE and transferred to PVDF membranes, which were probed with the indicated primary antibodies followed by fluorescently labelled secondary antibodies. Membranes were imaged using an Odyssey infrared imaging system under non-saturating conditions. Band intensities were quantified using Odyssey software with local background subtraction. Target protein levels were normalized to tubulin or to the appropriate fraction-specific marker, as indicated, and expressed relative to control samples. Data represent the mean ± SEM from at least three independent biological replicates.

### Nano-LC Mass Spectrometry

The gel bands were subjected to in-gel tryptic or AspN digestion using a DigestPro automated digestion unit (Intavis Ltd.) The resulting peptides were fractionated using an Ultimate 3000 nano-LC system in line with an Orbitrap Fusion Tribrid mass spectrometer (Thermo Scientific). In brief, peptides in 1% (vol/vol) formic acid were injected onto an Acclaim PepMap C18 nano-trap column (Thermo Scientific). After washing with 0.5% (vol/vol) acetonitrile 0.1% (vol/vol) formic acid peptides were resolved on a 250 mm × 75 μm Acclaim PepMap C18 reverse phase analytical column (Thermo Scientific) over a 150 min organic gradient, using 7 gradient segments (1-6% solvent B over 1min., 6-15% B over 58min., 15-32%B over 58min., 32-40%B over 5min., 40-90%B over 1min., held at 90%B for 6min and then reduced to 1%B over 1min.) with a flow rate of 300 nl min−1. Solvent A was 0.1% formic acid and Solvent B was aqueous 80% acetonitrile in 0.1% formic acid. Peptides were ionized by nano-electrospray ionization at 2.2 kV using a stainless-steel emitter with an internal diameter of 30 μm (Thermo Scientific) and a capillary temperature of 275°C.

All spectra were acquired using an Orbitrap Fusion Tribrid mass spectrometer controlled by Xcalibur 2.1 software (Thermo Scientific) and operated in data-dependent acquisition mode. FTMS1 spectra were collected at a resolution of 120 000 over a scan range (m/z) of 350-1550, with an automatic gain control (AGC) target of 400 000 and a max injection time of 100ms. Precursors were filtered according to charge state (to include charge states 2-7), with monoisotopic peak determination set to peptide and using an intensity range from 5E3 to 1E20. Previously interrogated precursors were excluded using a dynamic window (40s +/-10ppm). The MS2 precursors were isolated with a quadrupole mass filter set to a width of 1.6m/z. ITMS2 spectra were collected with an AGC target of 5000, max injection time of 50ms and HCD collision energy of 35%.

The raw data files were processed and quantified using Proteome Discoverer software v2.1 (Thermo Scientific) and searched against the UniProt Mus musculus database (downloaded April 2025; 54702 sequences) using the SEQUEST HT algorithm. Peptide precursor mass tolerance was set at 10ppm, and MS/MS tolerance was set at 0.6Da. Search criteria included oxidation of methionine (+15.995Da), acetylation of the protein N-terminus (+42.011Da), methionine loss plus acetylation of the protein N-terminus (−89.03Da) and phosphorylation of serine threonine and tyrosine (+79.966Da) as variable modifications and carbamidomethylation of cysteine (+57.021Da) as a fixed modification. Searches were performed with full tryptic or AspN digestion and a maximum of 2 missed cleavages were allowed. The reverse database search option was enabled and all data was filtered to satisfy false discovery rate (FDR) of 1%.

Label-free quantification of phosphopeptides was performed using peak area intensities obtained from Proteome Discoverer v2.1. Raw intensity values were log2-transformed prior to statistical analysis. Differential protein abundance analysis was performed using univariate paired t-tests. For all comparisons, the p-value was adjusted using the Benjamini-Hochberg FDR method. Volcano plots were generated by plotting log2 fold change against the negative log10 of the adjusted p-values. Phosphopeptides were considered significantly regulated if they met the criteria of an adjusted p-value (FDR) ≤ 0.05 and log2 fold change < −2.

## Supplementary Figures and Legends

**Figure S1.**
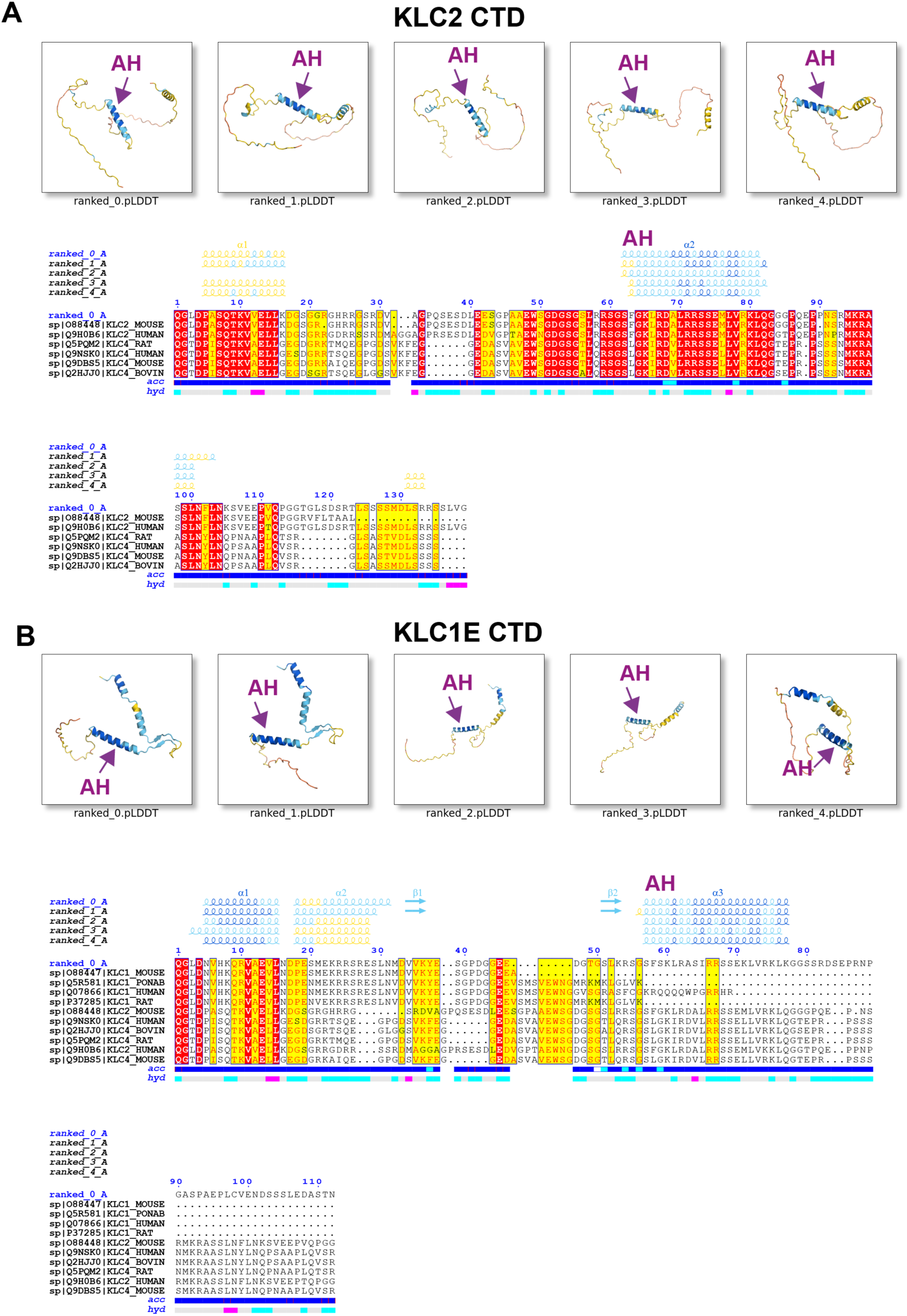
AlphaFold2 and FoldScript Analysis of KLC2 and KLC1E CTD. Structures CTD sequences for **(A)** KLC2 and **(B)** KLC1E (following the end of the structurally defined KLC TPR domain) to the carboxy terminus of the protein were predicted using AF2 and re-ranked using FoldScript. Models are coloured for pLDDT using standard AF2 scheme and alignments are annotated with predicted secondary structural features directly from FoldScript. Sequences used were KLC2 (NP_001356289.1 and KLC1E (NP_001020532.2) CTDs. In each case, the amphipathic helix (AH) is highlighted.

**Figure S2.**
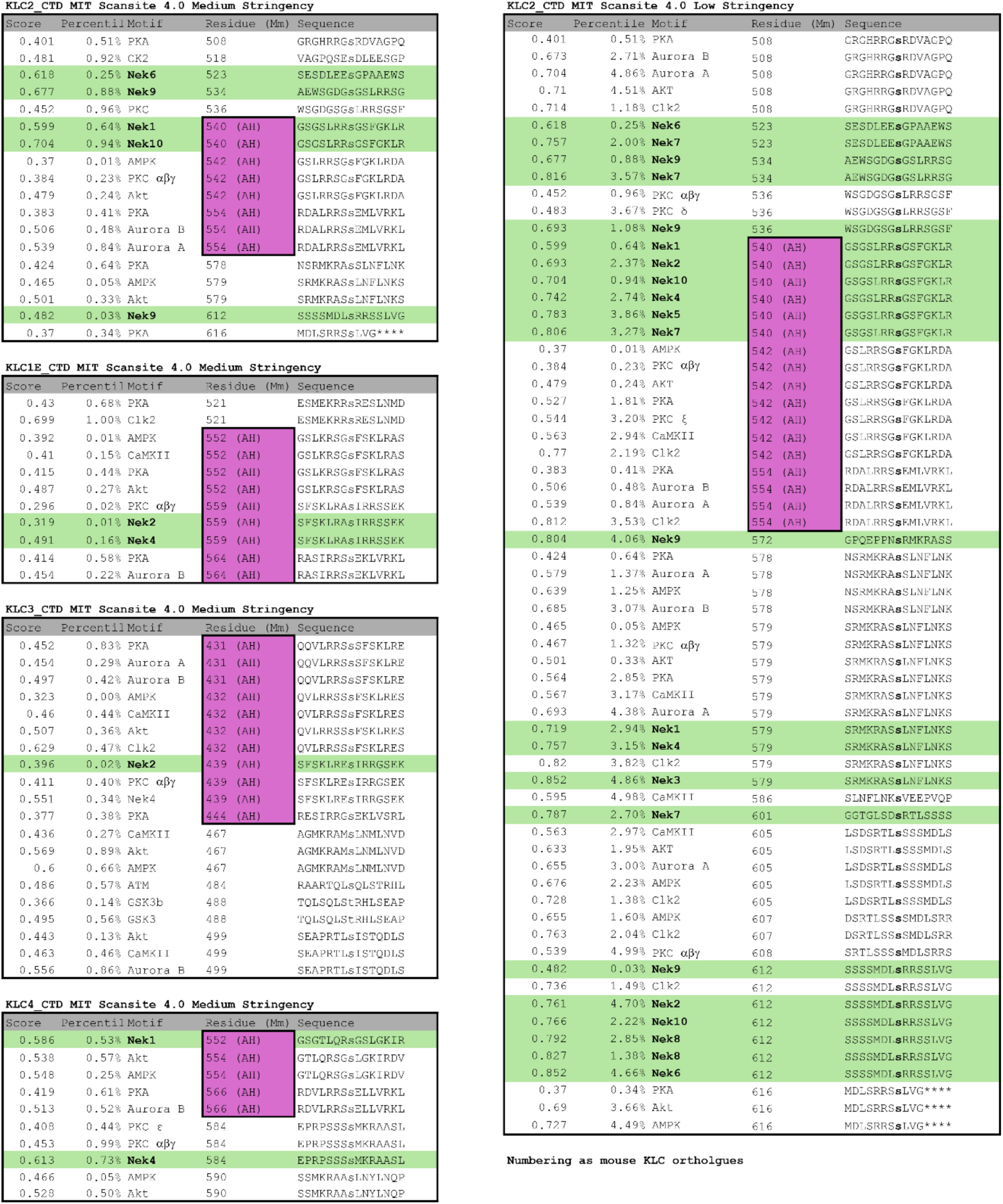
Scansite analysis of kinesin light chain carboxy-terminal domains. CTD sequences (following the end of the structurally defined KLC TPR domain) to the carboxy terminus of the protein were analysed used MIT Scansite 4.0 to identify motifs likely to be phosphorylated by specific protein kinases. Left shows medium stringency analysis for the mouse orthologues of KLC2 (NP_001356289.1), KLC1 isoform E (NP_001020532.2), KLC3 (NP_803136.2) and KLC4 (NP_001344059.1). Right shows low stringency analysis for KLC2. Predicted sites within the amphipathic helix are boxed in magenta. Potential NEK family kinase sites are highlighted in green. Numbering is as full-length protein. Scores and percentiles are as output by the software.

**Figure S3.**
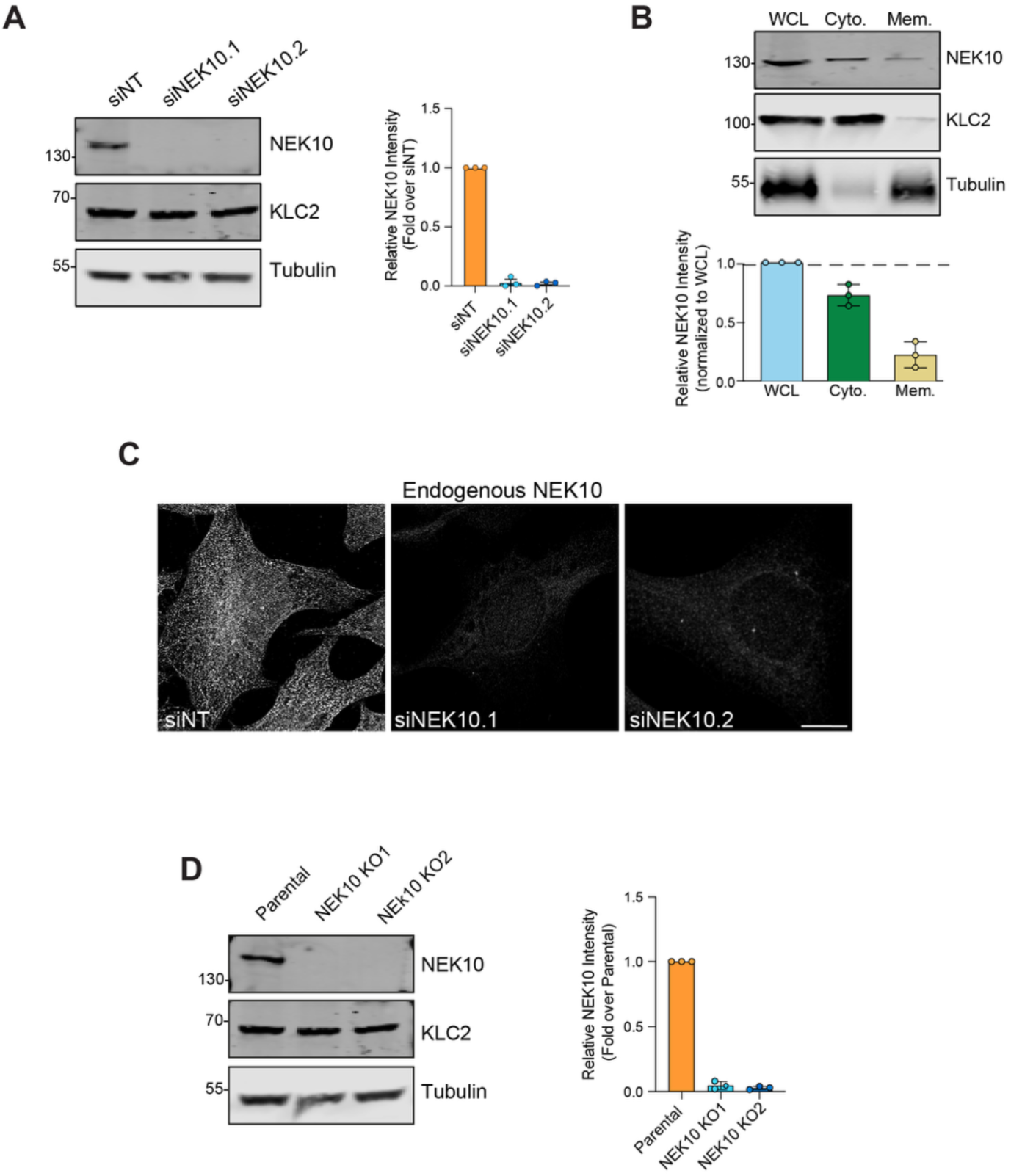
NEK10 siRNA knockdown and fractionation. (A) Representative western blot and quantification from three independent experiments showing NEK10 depletion using two different siRNAs (siNEK10.1, siNEK10.2). A non-targeting siRNA (siNT) was used as a negative control. (B) Western blot and quantification from three independent experiments showing results of cell fractionation experiment to examine relative NEK10 distribution. (C) Immunofluorescence imaging showing broadly distributed diffuse localisation for NEK10 and loss of signal following siRNA-mediated depletion. siNT, non-targeting siRNA. Scale bar is 10 µm. (D) Western blot and quantification from 3 separate experiments showing loss of endogenous NEK10 in NEK10 KO cell lines.

**Figure S4.**
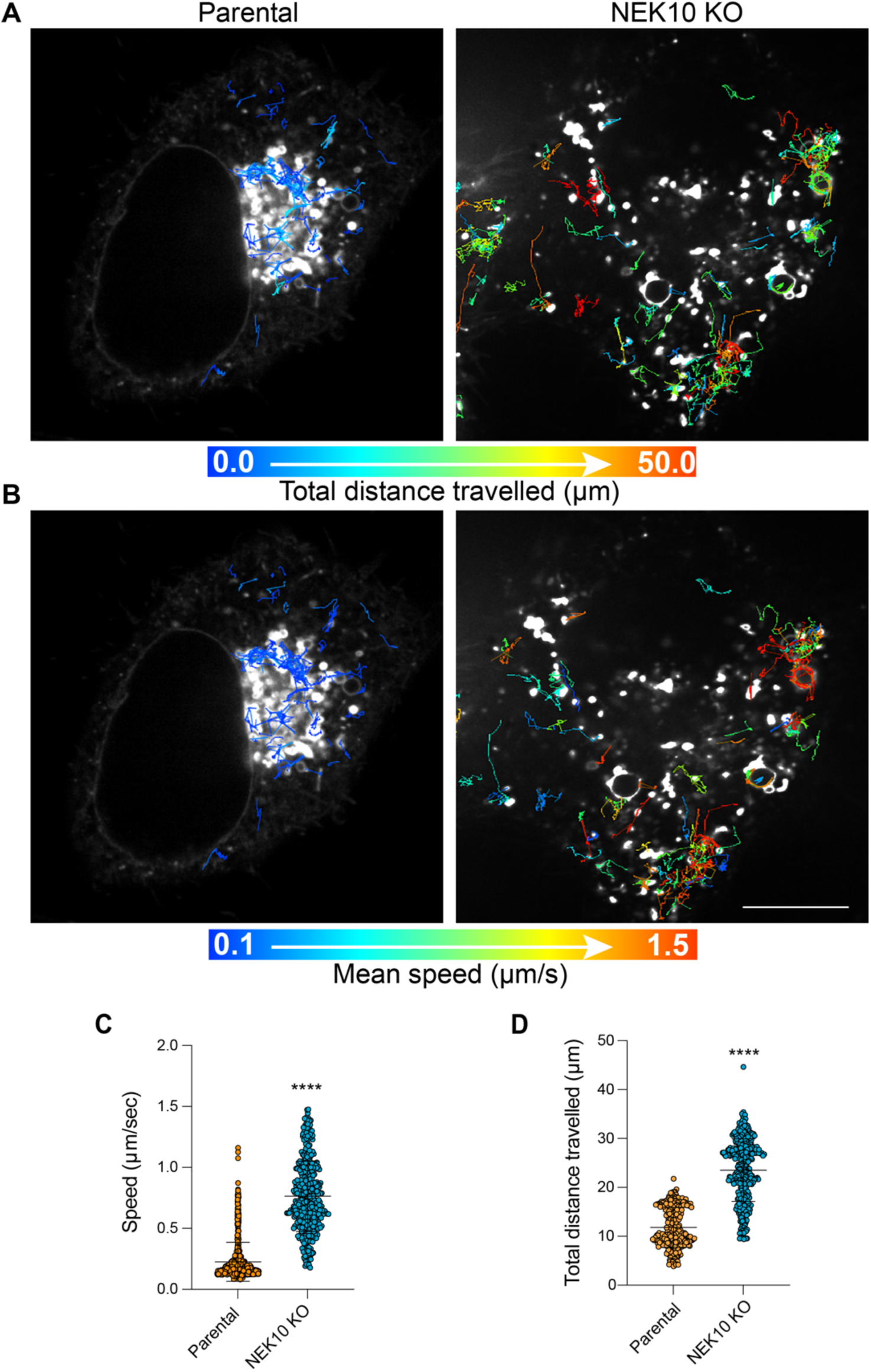
Analysis of lysosome motility in parental and NEK10 knockout Hela cells. Cells were transfected with LAMP-GFP and imaged live for 2 mins at 240ms per frame and trajectories were generated using TrackMate in Fiji. (**A**) & (**B**) Representative grayscale images showing first frame in sequence overlayed with lysosome tracks. Tracks are colour coded according to total distance travelled (**A**) or mean track speed (**B**). (**C**) & (**D**) Quantification of total distance travelled (C) or mean track speed (D) per lysosome track. Each point represents a single lysosome trajectory pooled from multiple cells and independent experiments. For statistical analysis, track values were averaged per cell and statical significance was determined using an unpaired two-tailed t-test on cell means. The bars represent Mean ± SD, where n = 3 independent experiments, with 10 cells analyzed per experiment. Scale Bar is 10 μm.

## Supplementary Movie Legends

**Movie S1. Spinning disk confocal imaging of LAMP1-GFP in parental Hela cells.** Images were acquired every 240 ms for 2 min. Scale Bar is 10 μm

**Movie S2. Spinning disk confocal imaging of LAMP1-GFP in NEK10 KO cells.** Images were acquired every 240 ms for 2 min. Scale Bar is 10 μm

## References

1. A. Yildiz, Mechanism and regulation of kinesin motors. Nat Rev Mol Cell Biol 26, 86–103 (2025).

2. J. A. Cross, M. P. Dodding, Motor-cargo adaptors at the organelle-cytoskeleton interface. Curr Opin Cell Biol 59, 16–23 (2019).

3. M. P. Dodding, M. Way, Coupling viruses to dynein and kinesin-1. Embo j 30, 3527–3539 (2011).

4. R. D. Vale, T. S. Reese, M. P. Sheetz, Identification of a novel force-generating protein, kinesin, involved in microtubule-based motility. Cell 42, 39–50 (1985).

5. S. Pernigo, A. Lamprecht, R. A. Steiner, M. P. Dodding, Structural basis for kinesin-1:cargo recognition. Science 340, 356–359 (2013).

6. Z. Antón et al., Molecular mechanism for kinesin-1 direct membrane recognition. Science Advances 7, eabg6636 (2021).

7. Y. Y. Yip et al., The light chains of kinesin-1 are autoinhibited. Proc Natl Acad Sci U S A 113, 2418–2423 (2016).

8. S. Pernigo et al., Structural basis for isoform-specific kinesin-1 recognition of Y-acidic cargo adaptors. eLife 7, e38362 (2018).

9. K. J. Verhey et al., Cargo of kinesin identified as JIP scaffolding proteins and associated signaling molecules. J Cell Biol 152, 959–970 (2001).

10. J. J. B. Cockburn et al., Insights into Kinesin-1 Activation from the Crystal Structure of KLC2 Bound to JIP3. Structure 26, 1486–1498.e1486 (2018).

11. M. P. Dodding, R. Mitter, A. C. Humphries, M. Way, A kinesin-1 binding motif in vaccinia virus that is widespread throughout the human genome. The EMBO Journal 30, 4523–4538 (2011).

12. S. Shukla et al. (eLife Sciences Publications, Ltd, 2025).

13. K. Chiba, K. M. Ori-McKenney, S. Niwa, R. J. McKenney, Synergistic autoinhibition and activation mechanisms control kinesin-1 motor activity. Cell Rep 39, 110900 (2022).

14. P. J. Hooikaas et al., MAP7 family proteins regulate kinesin-1 recruitment and activation. J Cell Biol 218, 1298–1318 (2019).

15. K. Barlan, W. Lu, V. I. Gelfand, The microtubule-binding protein ensconsin is an essential cofactor of kinesin-1. Curr Biol 23, 317–322 (2013).

16. A. E. McCart, D. Mahony, J. A. Rothnagel, Alternatively Spliced Products of the Human Kinesin Light Chain 1 (KNS2) Gene. Traffic 4, 576–580 (2003).

17. M. J. Woźniak, V. J. Allan, Cargo selection by specific kinesin light chain 1 isoforms. Embo j 25, 5457–5468 (2006).

18. T. Morihara et al., Transcriptome analysis of distinct mouse strains reveals kinesin light chain-1 splicing as an amyloid-β accumulation modifier. Proc Natl Acad Sci U S A 111, 2638–2643 (2014).

19. S. L. Chick et al., Whole-exome sequencing analysis identifies risk genes for schizophrenia. Nat Commun 16, 7102 (2025).

20. A. Gusev et al., Transcriptome-wide association study of schizophrenia and chromatin activity yields mechanistic disease insights. Nature Genetics 50, 538–548 (2018).

21. U. S. Melo et al., Overexpression of KLC2 due to a homozygous deletion in the non-coding region causes SPOAN syndrome. Hum Mol Genet 24, 6877–6885 (2015).

22. F. Bayrakli et al., Hereditary spastic paraplegia with recessive trait caused by mutation in KLC4 gene. Journal of Human Genetics 60, 763–768 (2015).

23. S. Kalantari, I. Filges, ’Kinesinopathies’: emerging role of the kinesin family member genes in birth defects. J Med Genet 57, 797–807 (2020).

24. C. Rosa-Ferreira, S. Munro, Arl8 and SKIP act together to link lysosomes to kinesin-1. Dev Cell 21, 1171–1178 (2011).

25. C. M. Guardia, G. G. Farías, R. Jia, J. Pu, J. S. Bonifacino, BORC Functions Upstream of Kinesins 1 and 3 to Coordinate Regional Movement of Lysosomes along Different Microtubule Tracks. Cell Rep 17, 1950–1961 (2016).

26. J. Pu, C. M. Guardia, T. Keren-Kaplan, J. S. Bonifacino, Mechanisms and functions of lysosome positioning. J Cell Sci 129, 4329–4339 (2016).

27. P. V. Hornbeck et al., PhosphoSitePlus, 2014: mutations, PTMs and recalibrations. Nucleic Acids Res 43, D512–520 (2015).

28. J. Jumper et al., Highly accurate protein structure prediction with AlphaFold. Nature 596, 583–589 (2021).

29. X. Robert, C. Guillon, P. Gouet, FoldScript: a web server for the efficient analysis of AI-generated 3D protein models. Nucleic Acids Research 53, W277–W282 (2025).

30. N. van Hilten et al., PMIpred: a physics-informed web server for quantitative protein– membrane interaction prediction. Bioinformatics 40, (2024).

31. J. C. Obenauer, L. C. Cantley, M. B. Yaffe, Scansite 2.0: Proteome-wide prediction of cell signaling interactions using short sequence motifs. Nucleic Acids Res 31, 3635–3641 (2003).

32. B. van de Kooij et al., Comprehensive substrate specificity profiling of the human Nek kinome reveals unexpected signaling outputs. eLife 8, e44635 (2019).

33. P. Dutt, N. Haider, S. Mouaaz, L. Podmore, V. Stambolic, β-catenin turnover is regulated by Nek10-mediated tyrosine phosphorylation in A549 lung adenocarcinoma cells. Proceedings of the National Academy of Sciences 121, e2300606121 (2024).

34. R. R. Chivukula et al., A human ciliopathy reveals essential functions for NEK10 in airway mucociliary clearance. Nat Med 26, 244–251 (2020).

35. A. Peres de Oliveira et al., NEK10 interactome and depletion reveal new roles in mitochondria. Proteome Science 18, 4 (2020).

36. N. Haider et al., NEK10 tyrosine phosphorylates p53 and controls its transcriptional activity. Oncogene 39, 5252–5266 (2020).

37. L. S. Moniz, V. Stambolic, Nek10 mediates G2/M cell cycle arrest and MEK autoactivation in response to UV irradiation. Mol Cell Biol 31, 30–42 (2011).

38. K. M. Power et al., NEKL-4 regulates microtubule stability and mitochondrial health in ciliated neurons. Journal of Cell Biology 223, (2024).

39. J. F. Weijman et al., Molecular architecture of the autoinhibited kinesin-1 lambda particle. Science Advances 8, eabp9660 (2022).

40. J. A. Cross et al., A de novo designed coiled coil-based switch regulates the microtubule motor kinesin-1. Nat Chem Biol 20, 916–923 (2024).

41. B. Cabukusta, J. Neefjes, Mechanisms of lysosomal positioning and movement. Traffic 19, 761–769 (2018).

42. J. A. Cross, M. S. Chegkazi, R. A. Steiner, D. N. Woolfson, M. P. Dodding, Fragment-linking peptide design yields a high-affinity ligand for microtubule-based transport. Cell Chem Biol 28, 1347–1355.e1345 (2021).

43. A. M. Fry, R. Bayliss, J. Roig, Mitotic Regulation by NEK Kinase Networks. Front Cell Dev Biol 5, 102 (2017).

44. A. M. Fry, L. O’Regan, S. R. Sabir, R. Bayliss, Cell cycle regulation by the NEK family of protein kinases. Journal of Cell Science 125, 4423–4433 (2012).

45. G. Chen et al., Kinesin-2 autoinhibition requires elbow phosphorylation. eLife 13, RP103648 (2025).

46. S. N. Cullati, L. Kabeche, A. N. Kettenbach, S. A. Gerber, A bifurcated signaling cascade of NIMA-related kinases controls distinct kinesins in anaphase. Journal of Cell Biology 216, 2339–2354 (2017).

47. A. J. Faragher, A. M. Fry, Nek2A kinase stimulates centrosome disjunction and is required for formation of bipolar mitotic spindles. Mol Biol Cell 14, 2876–2889 (2003).

48. J. Rapley et al., The NIMA-family kinase Nek6 phosphorylates the kinesin Eg5 at a novel site necessary for mitotic spindle formation. J Cell Sci 121, 3912–3921 (2008).

49. J. R. Mann et al., Loss of function of the ALS-associated NEK1 kinase disrupts microtubule homeostasis and nuclear import. Science Advances 9, eadi5548 (2023).

50. S. Cohen, A. Aizer, Y. Shav-Tal, A. Yanai, B. Motro, Nek7 kinase accelerates microtubule dynamic instability. Biochim Biophys Acta 1833, 1104–1113 (2013).

51. R. Adib et al., Mitotic phosphorylation by NEK6 and NEK7 reduces the microtubule affinity of EML4 to promote chromosome congression. Science Signaling 12, eaaw2939 (2019).

52. T. Ichimura et al., Phosphorylation-dependent interaction of kinesin light chain 2 and the 14-3-3 protein. Biochemistry 41, 5566–5572 (2002).

53. E. D. McKenna, S. L. Sarbanes, S. W. Cummings, A. Roll-Mecak, The Tubulin Code, from Molecules to Health and Disease. Annu Rev Cell Dev Biol 39, 331–361 (2023).

54. M. Mirdita et al., ColabFold: making protein folding accessible to all. Nat Methods 19, 679–682 (2022).

55. E. F. Pettersen et al., UCSF Chimera--a visualization system for exploratory research and analysis. J Comput Chem 25, 1605–1612 (2004).

